## Supplemental table1; supplemental figure 1 for "DNA metabarcoding reveals the dietary profiles of a benthic marine crustacean, *Nephrops norvegicus*"

Supplementary tables

**Table S1** Dietary overlap using Horn's Similarity Index among sites.

| | East | Irish | West |
|---|---|---|---|
| East NS | | 0.000382 | 0.030763 |
| Irish S | | | 0.029665 |
| West NS | | | |

Supplementary figures

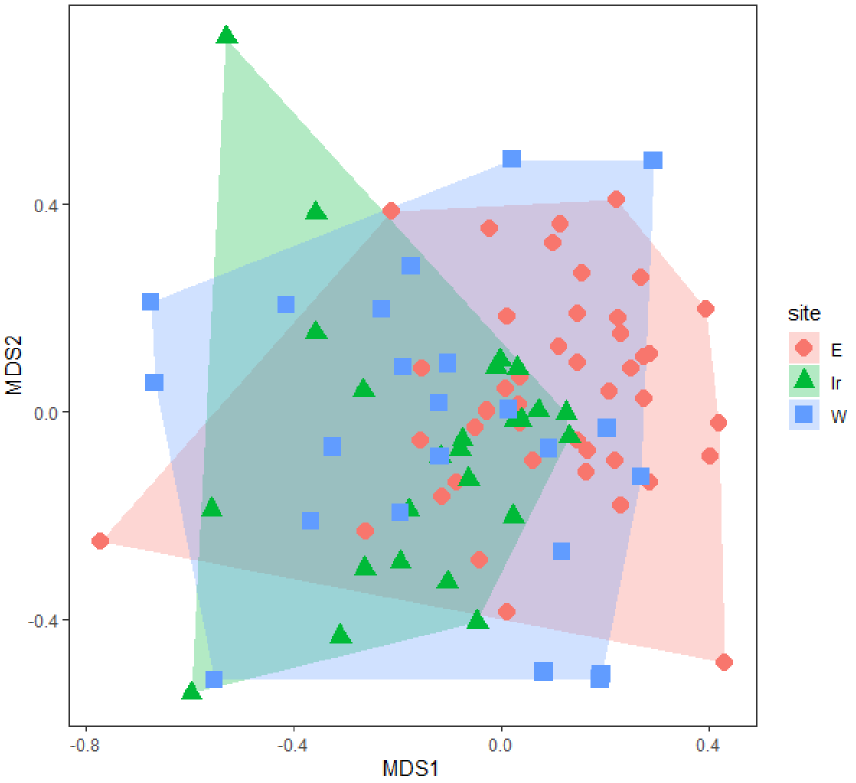


**Figure S1** Multidimensional Scaling (MDS) plot of diet variation of *Nephrops norvegicus* between sites, E: East (North Sea); Ir: Irish Sea; W: West (North Sea).

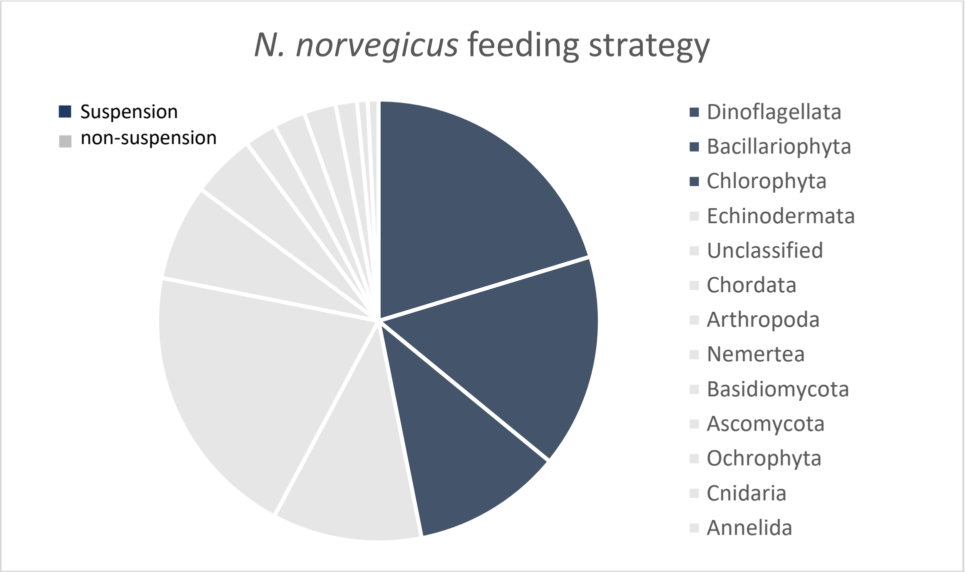


**Figure S2** Pie chart illustrating *N. norvegicus* feeding strategy. Food likely consumed through suspension feeding are indicated by blue and non-suspension indicated by grey.
